# Gross anatomy of the Pacific hagfish, *Eptatretus burgeri*, with special reference to the coelomic viscera

**DOI:** 10.1101/2022.12.08.519682

**Authors:** Banri Muramatsu, Daichi G. Suzuki, Masakazu Suzuki, Hiroki Higashiyama

## Abstract

Hagfish (Myxinoidea) are a deep-sea taxon of cyclostomes, the extant jawless vertebrates. Many researchers have examined the anatomy and embryology of hagfish to shed light on the early evolution of vertebrates; however, the diversity within hagfish is often overlooked. Hagfish have two lineages, Myxininae and Eptatretinae. Usually, textbook illustrations of hagfish anatomy reflect the morphology of the former lineage, especially *Myxine glutinosa*, with its single pair of external branchial pores. Here, we instead report the gross anatomy of an Eptatretinae, *Eptatretus burgeri*, which has six pairs of branchial pores, especially focusing on the coelomic organs. Dissections were performed on fixed and unfixed specimens to provide a guide for those doing organ- or tissue-specific molecular experiments. Our dissections revealed that the ventral aorta is Y-branched in *E. burgeri*, which differs from the unbranched morphology of *Myxine*. Otherwise, there were no differences in the morphology of the lingual apparatus or heart in the pharyngeal domain. The thyroid follicles were scattered around the ventral aorta, as has been reported for adult lampreys. The hepatobiliary system more closely resembled those of jawed vertebrates than those of adult lampreys, with the liver having two lobes and a bile duct connecting the gallbladder to each lobe. Overall, the visceral morphology of *E. burgeri* does not differ significantly from that of the known *Myxine* at the level of gross anatomy, except for the number of branchial pores.

## INTRODUCTION

One key to understanding the early evolution of vertebrates is to examine the morphology of our jawless sister group, the cyclostomes. This group and the gnathostomes (jawed vertebrate lineage) diverged about 500 million years ago (Kuraku & Kuratani, 2006). The cyclostome lineage has two subgroups: the lampreys (Petromyzontiformes) and hagfishes (Myxinoidea). Historically, these taxa have sometimes been considered paraphyletic, but today their monophyly is supported by molecular phylogeny (Mallatt & Sullivan, 1998; Kuraku et al., 1999; Takezaki et al., 2003; Kuraku, 2008; Heimberg et al., 2010) and comparative morphology (Yalden, 1985; Ota et al., 2007; Oisi et al., 2013). The absence of many characteristics of jawed vertebrates (e.g., jaws, paired nostrils, limbs, cucullaris muscles, and vertebral centra) makes cyclostomes essential organisms for studying vertebrate evolution (Janvier, 1996; Ota et al., 2011). In addition, they have many cyclostome-specific features (e.g., lingual apparatus, velum, and mucous cartilage) not found in jawed vertebrates (Janvier, 1996; Oisi et al., 2013), and whether these features are inherited from the common ancestors of vertebrates or are synapomorphies of cyclostomes remains controversial (Yokoyama et al., 2021; also see Sugahara, 2021). The morphology of the hagfishes is regarded to be highly derived, including the degeneration of eyes because of adaptation to the deep sea (Gabbott et al., 2016; Dong & Allison, 2021), and many studies have instead used lampreys as a model for cyclostomes. However, recent studies have suggested that some lamprey characteristics that had been regarded as ancestral to the cyclostomes, such as the larval-type oral apparatus (Miyashita et al., 2021) and the transformation of the larval endostyle into the thyroid after metamorphosis (Takagi et al., 2022), are in fact the derived state from the acquisition of the ammocoete larval stage. Lampreys and hagfish split around 470 to 390 million years ago (Kuraku & Kuratani, 2006), and their morphology subsequently diverged. Thus, there is a strong need to gain a better understanding of the anatomy of hagfishes.

Although the morphology of the hagfishes is often thought to be similar among species, they are, in fact, quite diverse. Hagfish are comprised of 2 subfamilies (Myxininae and Eptatreninae), 6 genera, and 88 species worldwide (Fricke et al., 2022). Differences in their morphology are most apparent in their branchial pores and horny teeth, which are often used to identify species (Jørgensen et al., 1998; Nelson et al., 2016; Fig. 1). The external branchial pores are located in the middle part of the body (Fig. 1a). The number and morphology of these pores differ among hagfish species and are essential traits for taxonomy (Dean, 1904; Fig. 1b). For example, the Eptatretinae have more than five pairs of external branchial pores, whereas the Myxininae have a single pair (Nelson et al., 2016; Fig. 1b). Currently, much of the anatomical literature focuses on one Myxininae species, *Myxine glutinosa*, and one Eptatretinae species, *Eptatretus stoutii* (syn. *Bdellostoma stoutii)* (Müller, 1834; Dean, 1904; Cole, 1907; Marinelli & Strenger, 1956; Ziermann et al., 2014). Of these, the most frequently cited study is by Marinelli & Strenger (1956), who produced detailed anatomical drawings of *M. glutinosa*, which has a single pair of branchial pores.

**FIGURE 1.**
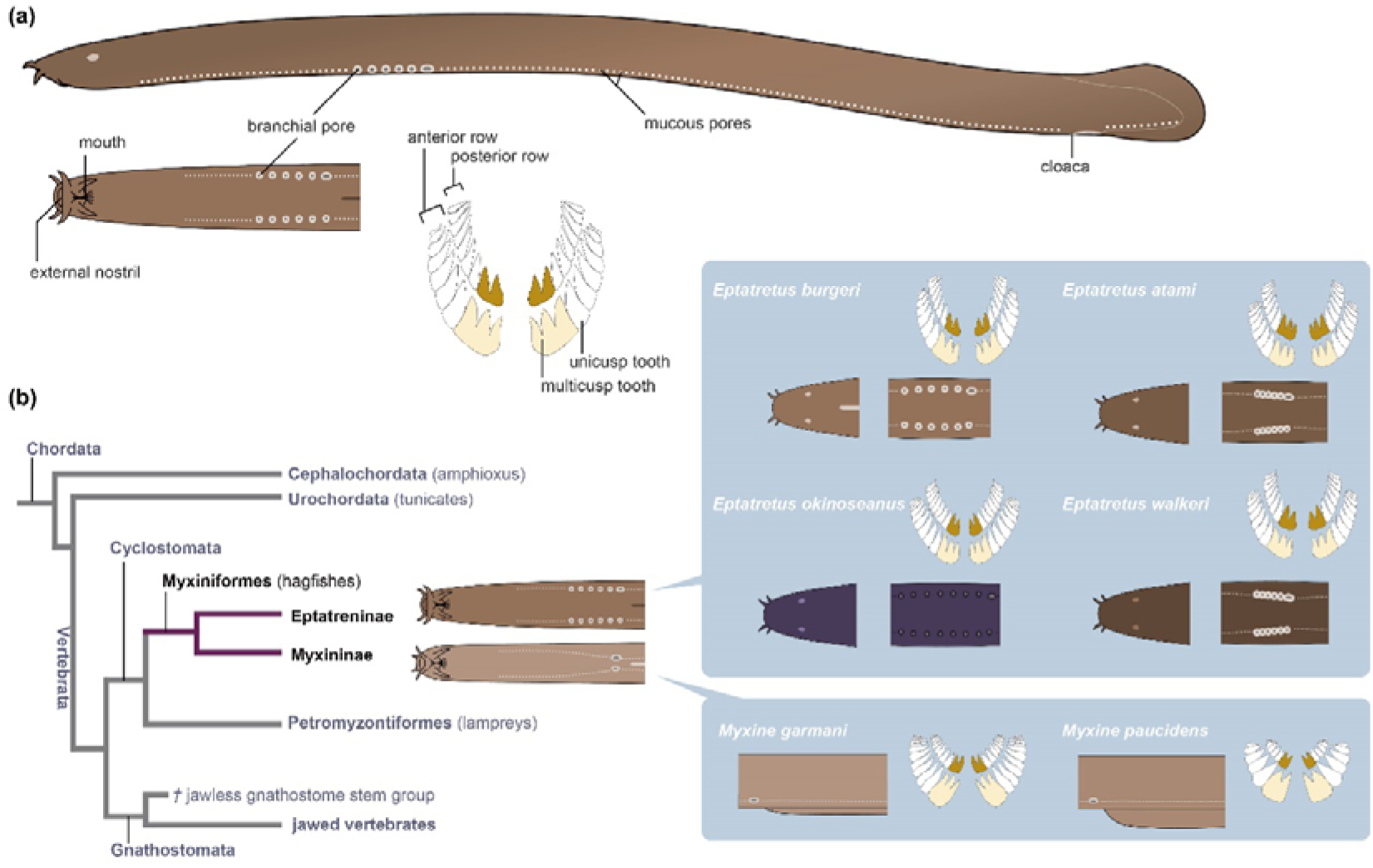
Morphological variation among hagfishes and their phylogenetic relationships. (a) Diagram of a hagfish, showing a left lateral view of the entire body, a ventral view of the head, and the tooth arrangement. (b) Phylogenetic relationships among the chordates. The Myxiniformes contain two lineages: Eptatreninae and Myxininae. The Eptatretinae have several pairs of external branchial pores, and the Myxininae have a single pair of branchial pores (Nelson et al., 2016). The morphological diagrams are after Nakabō (2013).

In the present paper, we focus on the anatomy of an Eptatretinae hagfish, the inshore hagfish *Eptatretus burgeri*. In East Asia, *E. burgeri* is commonly found in relatively shallow habitats. It is traditionally consumed and served as cuisine mainly in Korea and in some parts of Japan, where the species is easy to obtain (Gorbman et al., 1990; Honma, 1998). Recent embryological studies have been conducted on *E. burgeri*, and some knowledge on its comparative embryology is now available (Ota et al., 2007; Oisi et al., 2013, 2015). Gross anatomical description of a Japanese hagfish, probably this specie, has been done in the past in Japanese (Yamatsuta, 1903), but it is brief and not sufficient to discuss detailed comparative anatomy. Furthermore, we dissected unfixed specimens of hagfish, because we expect that further work will be conducted to examine the detailed molecular biology of this animal with techniques such as single-cell RNA sequencing. For this purpose, the tissue or organ of interest must be isolated appropriately through dissection; however, since fixation with formalin affects the preservation of RNA (Li et al., 2014; Denisenko et al., 2020), sample collection should be performed on unfixed specimens whenever possible. Because fixed tissues differ in appearance from unfixed ones, anatomical descriptions of fixed specimens may not be suitable for researchers who aim to conduct such studies. Thus, in this study, we compare *E. burgeri*, which has six pairs of branchial pores, to *M. glutinosa* to advance a comprehensive understanding of hagfish morphology and provide information on hagfish tissues using both unfixed and fixed specimens.

## MATERIALS AND METHODS

### Animals

*Eptatretus burgeri* specimens were collected from Sagami Bay, Kanagawa Prefecture, Japan, in February (40-45 m depth) and May (80-100 m depth) 2020. The animals from the former location were transported to the Center for Education and Research in Field Sciences, Faculty of Agriculture, Shizuoka University, Japan, and the latter were sent to the laboratory at the University of Tsukuba, Japan. Brown hagfish (*Eptatretus atami;* syn. *Paramyxine atami*) were obtained from Suruga Bay, Yaizu, Shizuoka Prefecture, Japan, in February 2021 (320 m depth), and transferred to the Field Science Center, Shizuoka University. The hagfishes were kept in seawater at 10 °C before dissection. We identified these species based on external morphology, including the branchial pores and dental cusps (after Dean, 1904; see Fig. 1). All animal experiments were performed in accordance with the Guides for Care and Use of Laboratory Animals of Shizuoka University and the University of Tsukuba.

### Anesthesia, fixation, and dissection

Hagfish were anesthetized with a mixture of 0.1% ethyl m-aminobenzoate (Nacalai tesque, Kyoto, Japan) and 0.1% sodium hydrogen carbonate (Wako, Osaka, Japan) in seawater. For fixation, we soaked the hagfish specimens in 20% formalin for two days and transferred them into 70% ethanol. We mainly followed Marinelli & Strenger (1956) for terminology.

## RESULTS

### External morphology

First, we describe the external morphology of *E. burgeri* (Fig. 2). *E. burgeri* is ochre to brownish red on its dorsal and lateral sides, with a pale dorsal midline skin fold, and white to ecru on its ventrum. Total lengths of our specimens were 39.5-58.0 cm. They have six pairs of external branchial pores (Fig. 2). The distance between each branchial pore is two or three times the branchial-pore diameter in *E. burgeri*. The caudal-most branchial pore on the left side is larger than the others because it is fused with the apertura ductus oesophageocutanei, as in other hagfish species (see also Dean, 1904).

**FIGURE 2.**
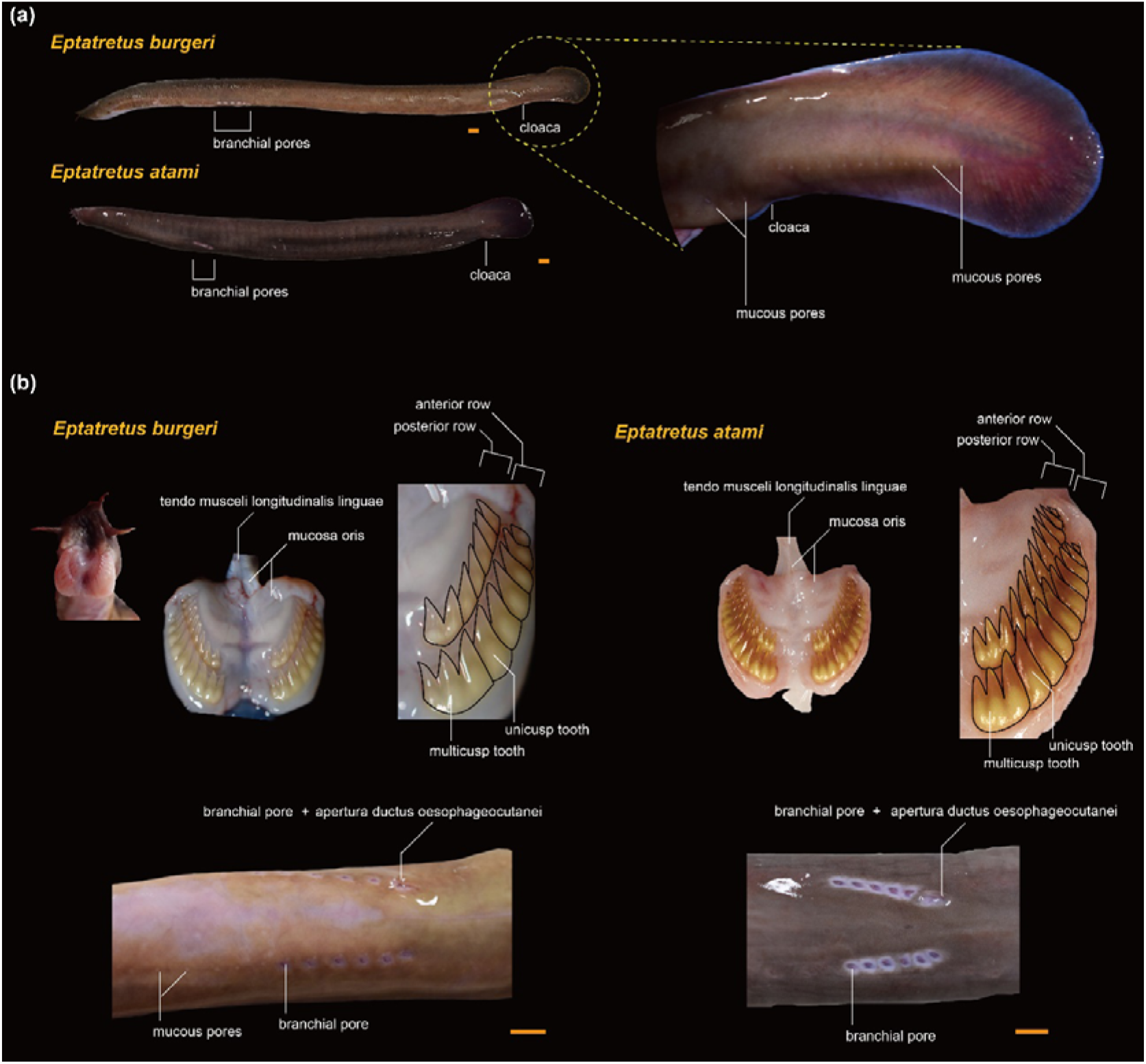
The external morphology of two hagfish species, *Eptatretus burgeri* and *E. atami*. (a) Photographs of each species, showing an enlarged view of the caudal part of *E. burgeri*. (b) Differences between the two species in the arrangement of tooth rows and external branchial pores. Scale bars = 1 cm.

Mucous pores open at regular intervals on the ventral side of the body in a pair of rows as with the external branchial pores. The rostral end of these rows is at about the same anterior–posterior position as the eyes, which is more anterior than the external branchial pores. These rows are also present on the tail, caudal to the cloaca, although they are interrupted around the cloaca (Fig. 2a). Rows of white and prolate-shaped mucous glands (glandular mucosa) are found under the mucous pores (Fig. 3).

**FIGURE 3.**
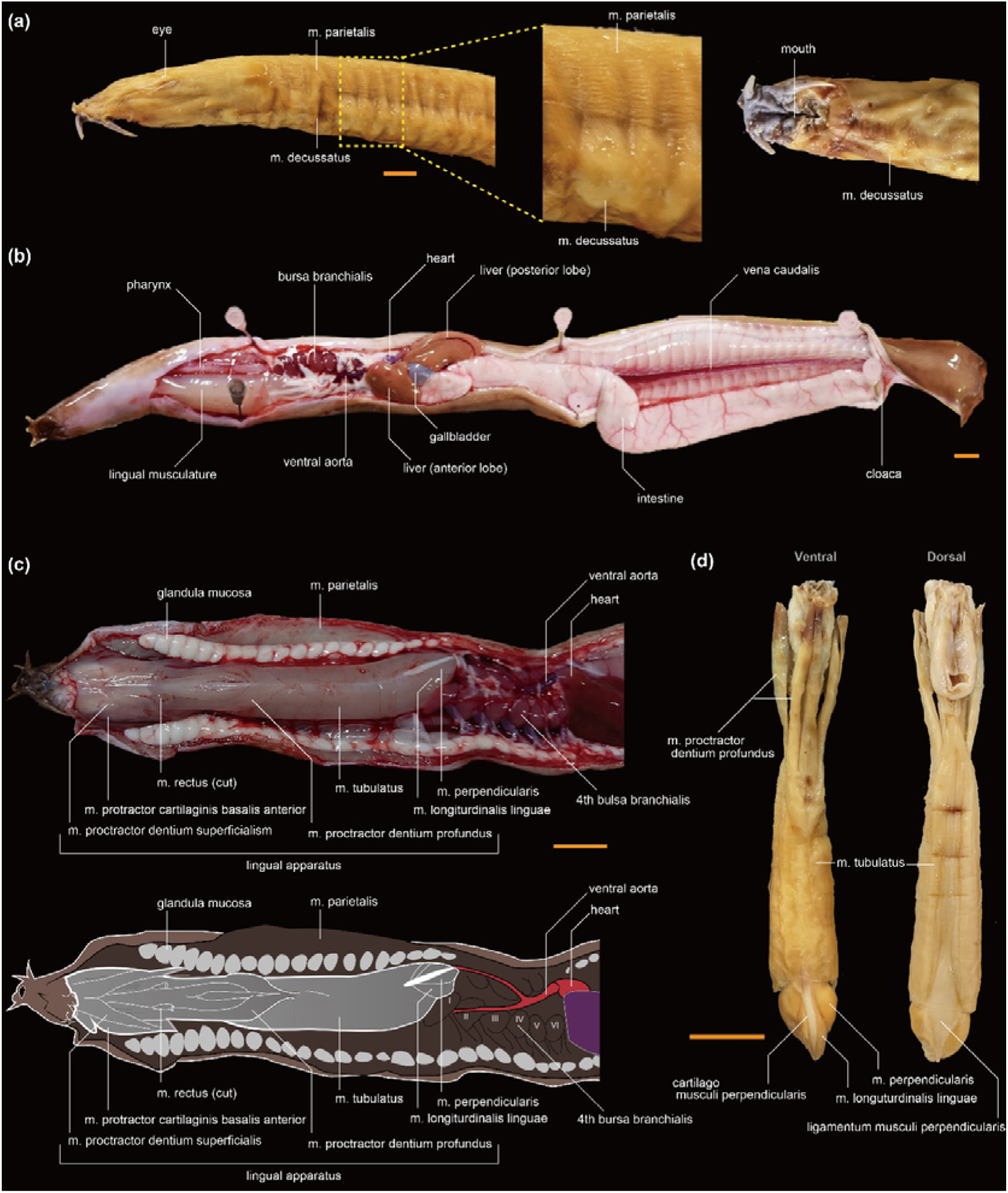
Positions of the organs and lingual apparatus of *Eptatretus burgeri*. (a) Fixed specimen with skin removed. The trunk is covered by two muscles with different fiber directions. (b) We cut and opened the body wall muscle using an unfixed specimen to show the lingual apparatus and internal organs. The photograph shows a ventral view. (c) Pharynx and lingual apparatus. (d) The lingual apparatus of a fixed specimen. Scale bars = 1 cm.

On the dental plate, *E. burgeri* has three fused cusps in the anterior row and two fused cusps in the posterior row (Fig. 2). The number of unicusp teeth varies among species, with some within-species differences (Jørgensen et al., 1998; Kase et al., 2017). Three pairs of tentacles are present on either side of the snout.

The above features are not so different in the other specie in the same genus, *E. atami* (Fig. 2). *E. atami* is dark brown or blackish purple; the total lengths were 43.6-55.3 cm. The notable differences in the number of dental casps and the distance between the branchial pores in *E. atami* compared to *E. burgeri*, but these two species have identical topographical relationships, including the internal organs. Thus, although there are minor differences in coloration and teeth, the number and topography of anatomical structures, such as branchial pores and tentacles, are highly conserved in the same *Eptatretus* genus.

### Lingual apparatus

The body trunk of hagfishes is broadly covered with a thin layer of m. parietalis and m. decussatus (Fig. 3a). For unfixed *E. burgeri* specimens, we pinched the ventral skin rostral to the branchial region and just ventral to the lingual apparatus with tweezers and opened the abdomen along the midline (Fig. 3b, c).

The lingual apparatus is unique to cyclostomes, and differs from the tongue (hypobranchial musculature) of jawed vertebrates in that it is derived from the mandibular arch (i.e., the first pharyngeal arch; Oisi et al., 2015, and references within). The lingual apparatus is a prominent structure in the hagfish pharynx and provides mobility to the dental plate (Clark & Summers, 2007). It consists of the protractor muscles: m. protractor cartilaginis, m. protractor dentium superficialis, and m. protractor dentium profundus (Fig. 3c, d) that originate in the medial cartilaginous keel (cartilago linguae basalis) of the lingual apparatus and terminate in the dental plate. The main body of the thick rod-shaped lingual apparatus is dorsal to these protractor muscles. The rod-shaped structure is entirely covered by m. tubulatus, with the muscle fibers oriented horizontal to the body axis and encircling the rod-shaped structure. The longitudinal muscles (m. longitudinalis linguae and m. perpendicularis) are found medial to the m. tubulatus. These originate in the cartalago musculi perpendicularis and terminate at the posteromedial part of the dental plate with the stiff tendon (tendo musculi longitudinalis linguae). The muscles that make up the lingual apparatus were stiffened in our fixed specimens, but white and soft in unfixed samples with a texture that was clearly distinguishable from that of the other trunk muscles (Fig. 3c).

### Branchial apparatus, heart, and thyroids

The gills and heart are located caudal to the lingual apparatus in hagfish. These pharyngeal elements initially develop anterior to the head in the hagfish embryo, as in other vertebrates. During development, the anterior half of the head becomes elongated, and the gills and heart are shifted posteriorly (Oisi et al., 2013).

The branchial apparatus of hagfish has a unique morphology: the branchial pouch forms a bladder-shaped structure (bursa branchialis) that connects with the pharynx and external body through ducts (Marinelli & Strenger, 1956). In *E. burgeri*, we observed six pairs of hemispherical bulsa branchialis lined up on each side (Fig. 4a). The inner gill duct (ductus branchialis afferens) connects the pharynx and bulsa branchialis. The outer gill duct (ductus branchialis efferens) emerges from the ventral side of the bulsa branchialis and is connected to the branchial pore (Fig. 4a).

**FIGURE 4.**
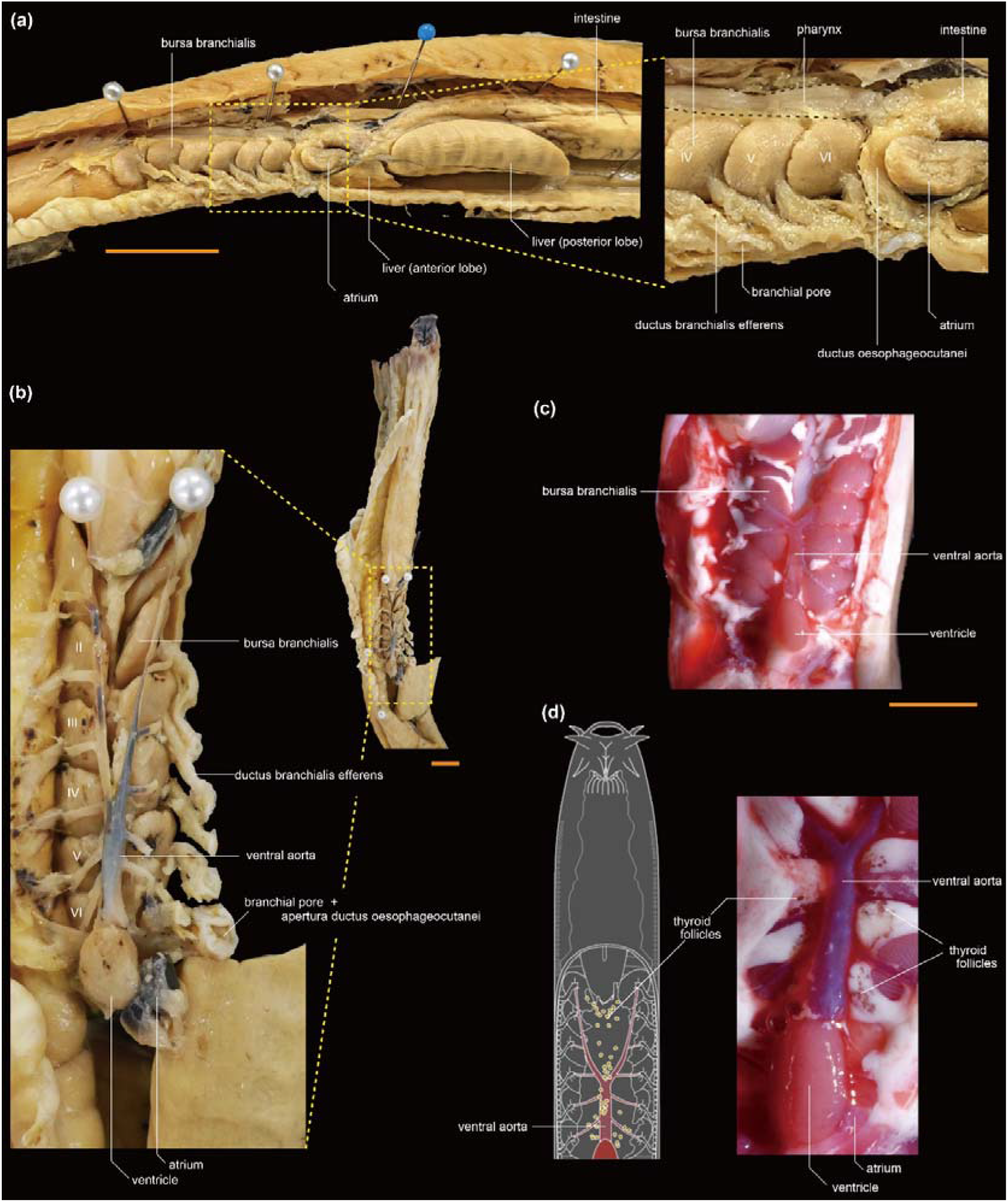
Cardiac anatomy and thyroid follicles of *Eptatretus burgeri*. (a) Left lateral view of a fixed specimen. We removed the lateral body wall and exposed the left lateral branchial structure. (b) Ventral view of the branchial organs of the same specimen as shown in panel (a). The left lateral body wall was removed, exposing the left-sided rows of the ductus branchialis efferens. (c) Ventral view of the branchial structure in an unfixed specimen. (d) Ventral view of the heart in an unfixed specimen. The thyroid follicles are in the white adipose tissue. The distribution of the thyroid follicles is shown in the diagram. Scale bars = 1 cm.

On the left side of the body, caudal to the caudal-most ductus branchialis efferens, we identified the ductus oesophageocutanei, a structure unique to hagfish (Fig. 4b). This duct directly connects the alimentary canal to the exterior of the body at the boundary between the pharynx and intestine and is located only on the left side. The alimentary canal is slightly constricted at the branching point of this duct.

The heart is located along the midline, caudal to the caudal-most bulsa branchialis and rostral to the liver (Fig. 4b). The heart consists of a ventricle and atrium. The atrium is located caudal to and on the left side of the ventricle. As in other vertebrates, the ventral aorta extends anteriorly from the ventricles and leads to the branchial arteries (arteria branchialis afferens) in each bursa branchialis. The ventral aorta bifurcates around the branching point of the fourth branchial arteries (Fig. 4b–d), as already reported in *Bdellostoma cirrhatum* (syn. *E. burgeri)* in Yamatsuta (1903). This differs from the description of *M. glutinosa*, in which the ventral aorta does not bifurcate and remains along the midline (Marinelli & Strenger, 1956), and from that of *E. cirrhatus*, in which the ventral aorta bifurcates close to the ventricle (Icardo et al., 2016a, b). The surface of the heart and ventral aorta lacks the vascular system (e.g., coronary or hypobranchial arteries) seen in many jawed vertebrates (for coronary arteries of jawed vertebrates, see Mizukami et al., 2022).

In unfixed samples (Fig. 4c), the bulsa branchialis was easily identifiable as a flexible gill structure containing small blood vessels. White fat is found around this branchial region, and the thyroid follicles are embedded in it (Fig. 4d). The thyroid consists of yellow, 100-500-μm long oval thyroid follicles scattered around the ventral aorta from the anterior part of the ventricle to the posterior end of the lingual muscle (Fig. 4d).

### Hepatobiliary and urogenital organs

The hepatobiliary organ is found posterior to the heart. The liver is divided into anterior and posterior lobes, with the gallbladder located between them (Fig. 5a). The bile duct, hepatic ducts, blood vessels, and connective tissues are also found between the two lobes and connect the hepatic organs with the gastrointestinal tract. In unfixed specimens, the gallbladder was a thin bladder filled with green bile (Fig. 5b). The surface of this bladder is covered with a network of small arteries. Apart from some large arteries, no vascular networks were visible on the gallblader in fixed specimens (Fig. 5c). Although the previous studie have suggested the presence of a pancreatic principal islet at the junction of the extrahepatic bile duct and intestine in hagfishes (Youson & Al-Mahrouki, 1999), we could not find this structure in the present gross dissection. Further histological investigation will be necessary to observe this pancreatic structure. Spleens were not found.

**FIGURE 5.**
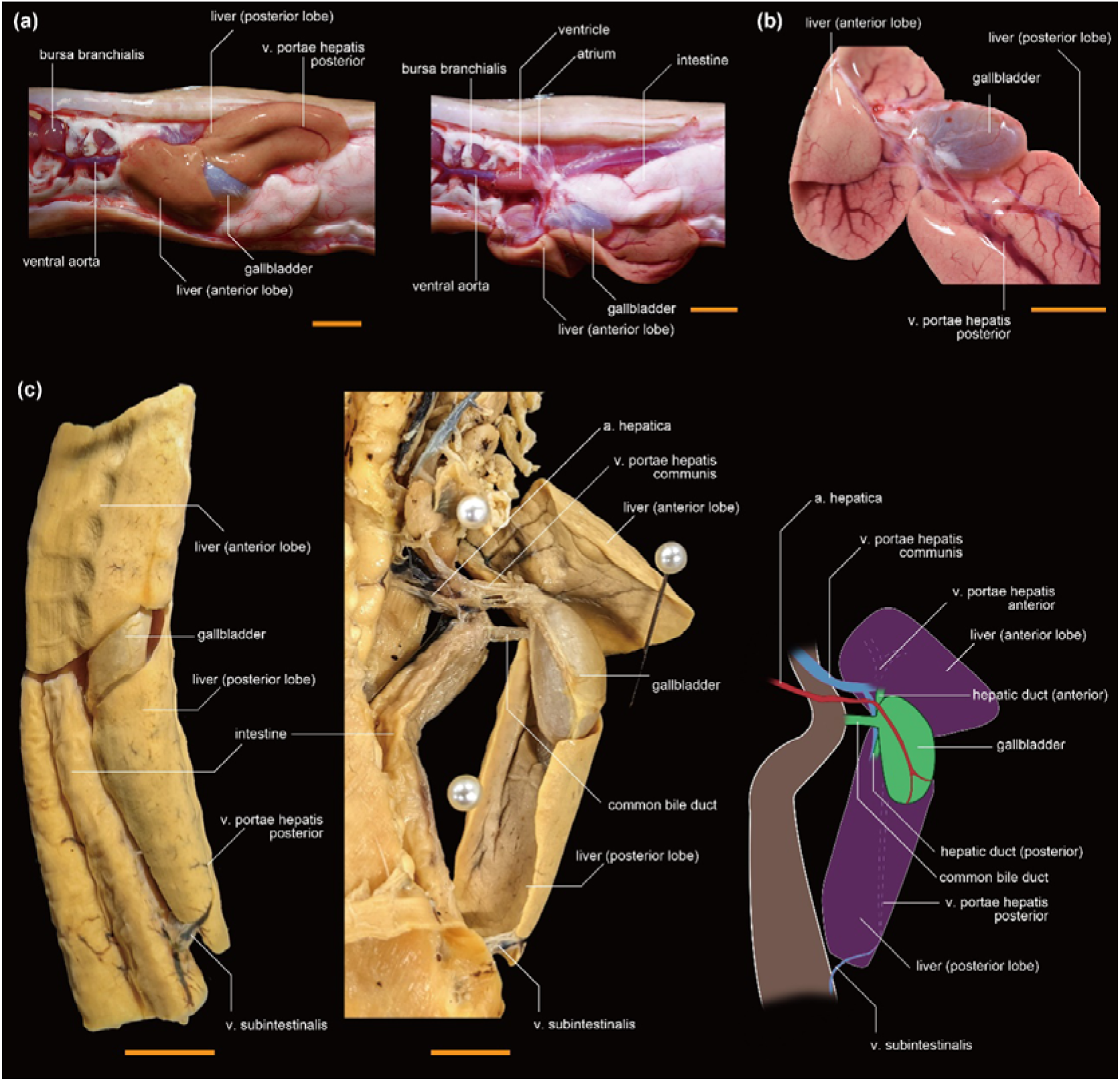
The hepatobiliary organs of *Eptatretus burgeri*. (a) Ventral views of an unfixed specimen. The liver in the left-hand panel is shown almost in its natural position. In the right-hand panel, we have shifted the liver to the left side of the body, showing the junction of the hepatobiliary system with the intestinal tract. (b) The hepatobiliary system of an unfixed specimen, where the liver and gallbladder were severed from the intestinal tract and are shown from the intestinal side. (c) The hepatobiliary system of a fixed specimen shown from the ventral side. The liver in the left-hand panel is in its natural position. The middle panel shows the liver flipped over to the right side of the body, showing the extrahepatic biliary tract and vascular system. Scale bars = 1 cm.

The common bile duct (ductus choledochus) is located between the two liver lobes, and transports bile from the gallbladder to the intestine (Fig. 5c). From the interface between the gallbladder and the common bile duct, two hepatic ducts branch off and enter the anterior and posterior lobes of the liver. These ducts are short, giving the appearance that the gallbladder is attached directly to the liver. The blood vessels on the gallbladder are the peripheral arteries branched from the coeliac artery (arteria coeliaca) that runs longitudinally along the intestinal tract. The portal vein (vena portae), a major vein in the liver, bifurcates proximal to the gallbladder and enters the anterior and posterior lobes of the liver from the same position as the anterior and posterior hepatic ducts. The posterior branch (v. portae hepatis posterior) passes near the ventral side of the posterior lobe of the liver. This vein passes out of the liver at the caudal edge of the posterior lobe and is distributed on the surface of the intestine, where it is termed the v. subintestinalis. We could not determine the peripheral pattern of v. portae hepatis anterior in the fixed specimens. In the unfixed specimens, we observed that this vessel branches all over the medial side (i.e., gallbladder side) of the anterior lobe of the liver (Fig. 5b).

The alimentary canal consists of a large simple intestine extending in a straight line (Fig. 6a). Unlike most jawed vertebrates, hagfish have no stomach. The intestinal tract of *E. burgeri* is a linear structure rich in blood vessels, as has been reported for other hagfishes. The intestinal wall is thick, elastic, and made up of longitudinally aligned folds on the inner surface (Fig. 6a).

**FIGURE 6.**
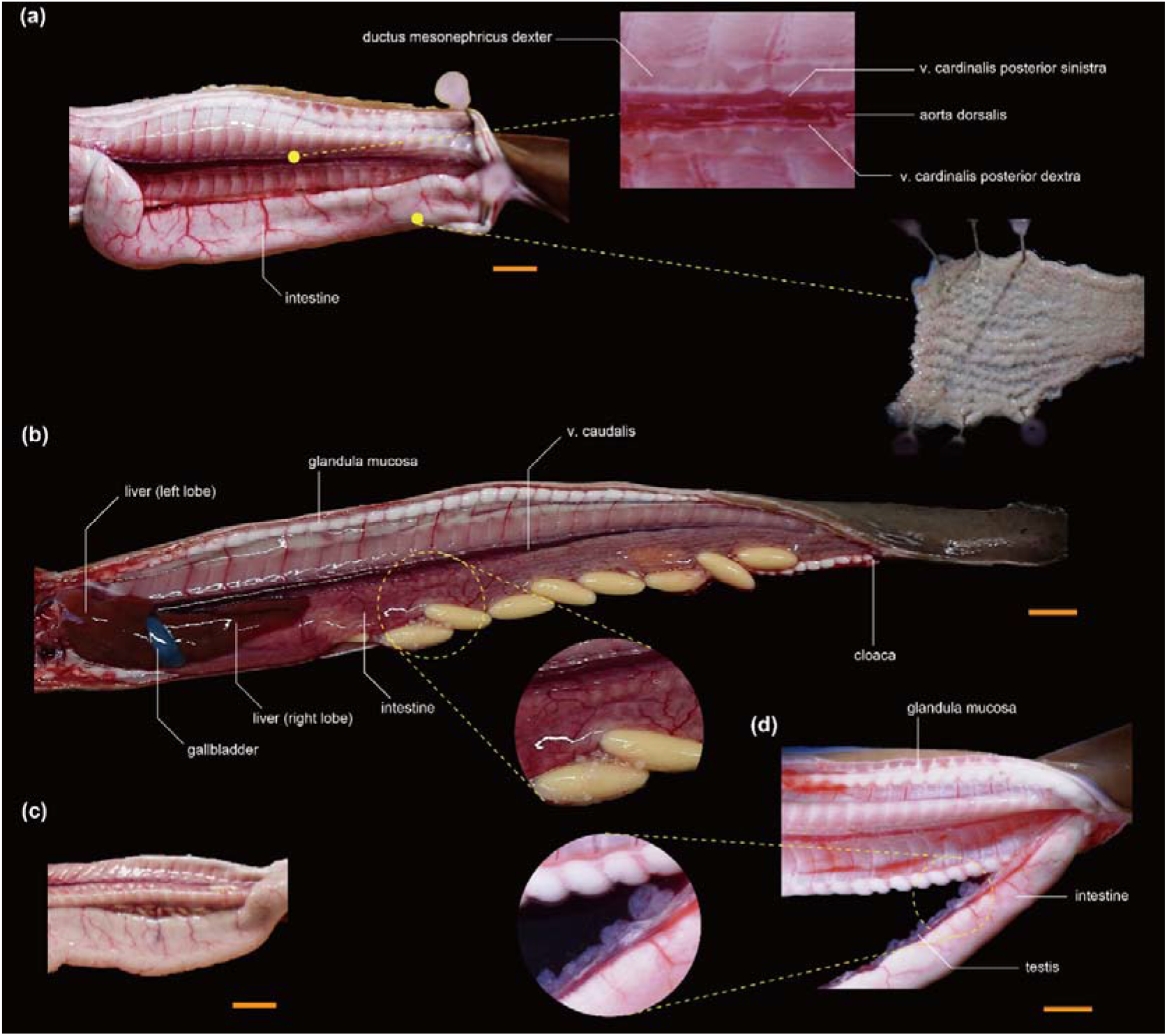
The hindgut and urogenital organs of unfixed specimens of *Eptatretus burgeri*. (a) The intestinal tract is shown from the ventral side. Enlarged views are shown of the kidneys and the inner wall of the intestinal tract. In this specimen, the gonads are not mature, and the sexes are not distinguishable. (b) The female body cavity with mature ovaries. (c) A female with immature ovaries. (d) A mature male specimen. Scale bars = 1 cm.

The kidneys are paired on the roof of the body cavity, as in other vertebrates, which are represented by the pronephros (see Romagnani et al., 2013). The gonads of hagfish are found in the peritoneal cavity and are attached to the intestinal tract. The testis consists of white-to cream-colored, translucent tissue (Fig. 6d), and the ovary is yellowish and opaque (Fig. 6b, c). In the waters around Japan, *E. burgeri* ovarian maturity is reported to occur in September, and the testes are at their maximum size around July (Patzner, 1978a, b; Nozaki et al., 2000). However, although we collected the hagfish in February, the gonads were in various stages of growth (Fig. 6b, c). The membranous ovary contains eggs at various stages of maturity. Immature eggs appear as small white prolate spheroids that turn yellow and oval as they mature. Sexual maturity can be confirmed in females by touching the abdomen from the outside.

### Head and brain

The m. parietaris continues serially from the trunk, covering the caudal half of the head, with the anterior end of this muscle reaching the posterior edge of the corbiculum nasale (Fig. 7a). The eyes are semi-transparent and easy to observe in the unfixed samples (Fig. 7a); in the fixed samples, their texture is different and the eyes are more difficult to identify (Fig. 3a). We needed to remove the m. parietalis when we observed the brain because this muscle covers the cranial roof. The dorsal surface of the brain is covered with thin cartilage, which must be removed with scissors for observation of the brain. There is little space between the brain and the brain capsule (Fig. 7a). The brain consists of the telencephalon, diencephalon, mesencephalon, and rhombencephalon (Fig. 7b; also see Dupret et al., 2014; Sugahara et al., 2017; Suzuki, 2021), as in other vertebrates. There is no overt epiphysis. The habenular ganglion (ganglion habenulae) is located on the midline between the telencephalon and the diencephalon. On each side of the rhombencephalon is a cartilaginous auditory capsule (capsula auditiva) with a single-canaled inner ear. This single canal is a derived condition among the cyclostomes (Higuchi et al., 2019). At the rostral end of the brain, the fila olfactoria enter the nasal cavity and is closely attached to the corbiculum nasale, making it difficult to detach without damaging it. Brain texture changed significantly after fixation. Because the brain loses flexibility and becomes brittle after fixation, isolation of the brain should be performed in the unfixed condition.

**FIGURE 7.**
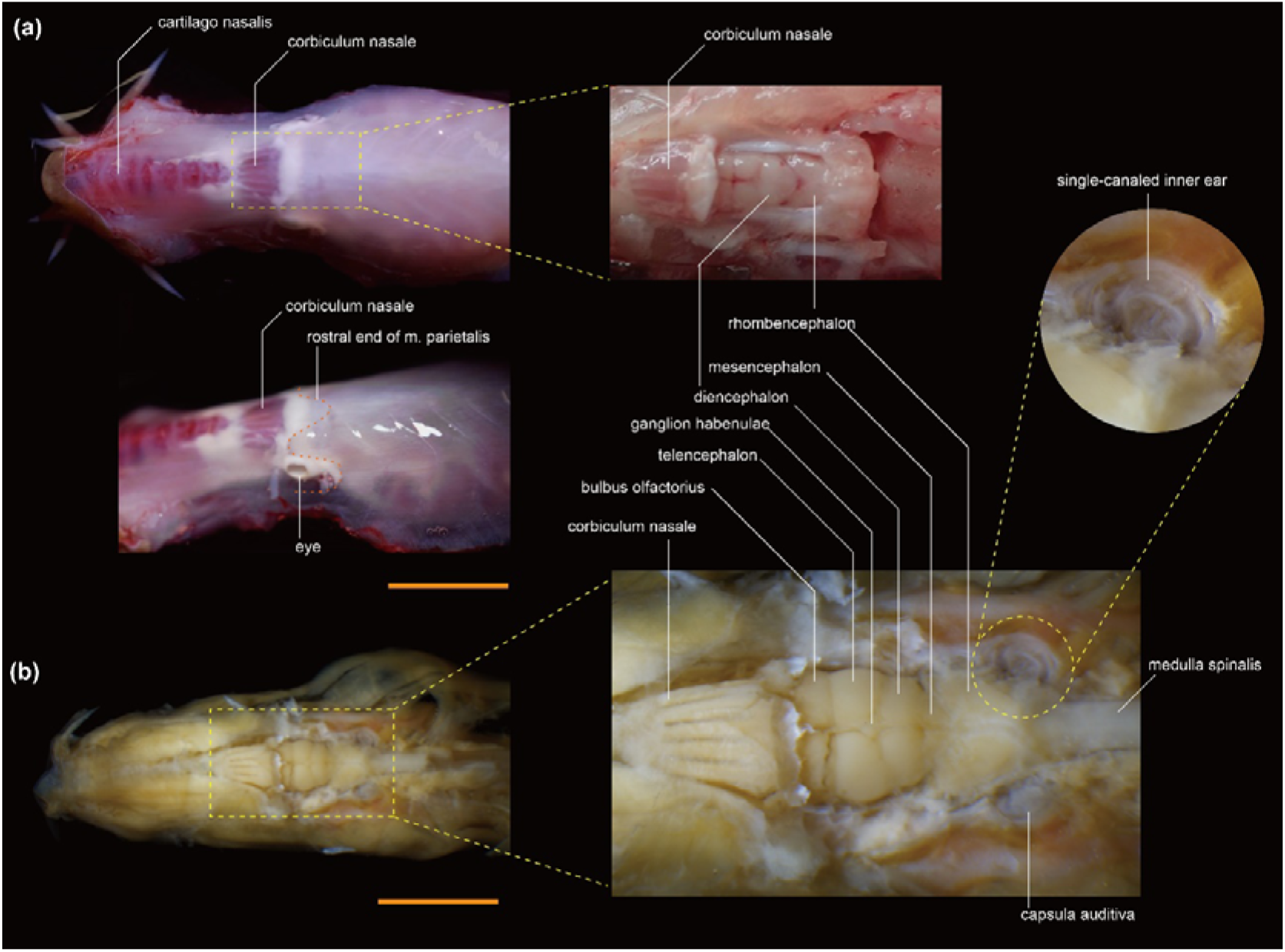
The head of *Eptatretus burgeri*, with the skin removed. (a) The unfixed specimen. (b) The fixed specimen. Scale bars = 1 cm.

## DISCUSSION

### Caudal shift of cardiobranchial structures and enlargement of the lingual apparatus

The division of the head and trunk appears to be ambiguous in adult hagfish. Their bodies are covered with segmented m. parietalis up to the area surrounding the brain, and the branchial pores open at about the middle of the body. This should be related to two developmental processes: the caudal shift of the pharyngeal structures and the rostral shift of the trunk somite derivatives, as observed in the development of *E. burgeri* (see Oisi et al., 2015).

The vertebrate heart is originally a structure closely associated with the pharyngeal arch, and the head mesoderm and cranial neural crest cells contribute to it during the pharyngula stage (Tzahor & Evans, 2011; Meilhac & Buckingham, 2018). Because of this, the heart is located in the head near the pharynx in most fishes, but adults of some tetrapod groups are the exception. A typical example is amniotes, in which the heart shifts caudally to the thorax during development, away from the cranium, to establish a long neck (Hirasawa et al., 2016). This translocation of the heart during the establishment of a long neck results in the heart being surrounded by trunk derivatives, such as the rib cage.

The caudal shift of the branchial and heart region in hagfishes differs from the translocation of the heart in amniotes. In jawed vertebrates, the mandibular arch differentiates into the jaw and the hyoid arch into the hyoid apparatus and cutaneous muscles of the neck. They are rarely remodeled, whether the neck is long or not. Remodeling occurs instead in the caudal-most part of the pharyngeal arch series, resulting in the extension of the esophagus behind the caudal-most pharynx, giving rise to a long neck. By contrast, in the hagfishes, elongation occurs in the rostral pharynx. The fourth pharyngeal pouches as well as more caudal ones differentiate as simple branchial pores and shift caudally with the heart following the remodeling of the first through third pharyngeal arches (Oisi et al., 2015). Thus, the anterior part of the pharynx, not the esophagus, is elongated in hagfishes without establishing a neck. Simultaneously, the trunk somites shift rostrally to cover the head surface (Oisi et al., 2015). This is also an important difference from jawed vertebrates, whose necks are characterized by cucullaris muscles (Oisi et al., 2015 and references therein).

This rostral extension of the somites is also found in lampreys (Kuratani et al., 1999), suggesting that the presence of trunk muscles surrounding the heart is common to all adult cyclostomes. In lampreys, however, the translocation of the cardiobranchial region does not occur. A large lingual apparatus occupies the space between the cranium and the branchial region in hagfishes (Fig. 3), whereas the lingual apparatus is smaller and located on the ventral side of the branchial region in lampreys (Marinelli & Strenger, 1954; Yalden, 1985). Thus, the shift in cardiobranchial position could be related to the enlargement of the lingual apparatus, a derivative of the mandibular arch. It is unclear which morphology, that of the lamprey or hagfish, more closely reflects the ancestral form of the cyclostomes. In *†Myxinikela siroka* and *†Tethymyxine tapirostrum*, which are regarded as close relatives of hagfish, the gills are much more cephalic in position than in the extant hagfishes (Bardack, 1991; Miyashita et al., 2019). Therefore, perhaps the enlargement of the lingual apparatus and the caudal shift of the cardiobranchial systems in the extant hagfishes is a derivative condition.

### Hagfish have unique diversities in the branchial region

*Myxine* and *Eptatretus* differ considerably in their external morphology in that the former group has a single pair of branchial pore apertures and the latter has multiple pairs. In *Myxine*, multiple ductus branchialis efferens open into a single branchial pore. In *Eptatretus*, each ductus branchialis efferens opens into a single branchial pore, which is thought to be the ancestral pattern (Jørgensen et al., 1998).

The number of bulsa branchialis and branchial pores varies throughout the Myxinoidean lineage. In *Eptatretus*, the number of bulsa branchialis has been reported to range from 5 to 14 pairs (Dean, 1904; Mincarone & McCosker, 2004). This not only varies among species but can also vary within a species. For example, in *Eptatretus stoutii* (syn. *Homea stouti)*, the number of gill pairs is usually 12 but can range from 10 to 15 (Dean, 1904). In *Myxine*, the number of bulsa branchialis does not vary as much as in *Eptatretus*, and ranges from five to seven pairs (Jørgensen et al. 1998). The cretaceous *†Tethymyxine tapirostrum* has eight pairs of bulsa branchialis (Miyashita et al., 2019).

In addition, perhaps related to the diversity of gill arch organs, there is also diversity in the branching pattern of the ventral aorta among hagfishes. In the *E. burgeri* specimens dissected in the present study, the ventral aorta branches in a “Y” pattern about a third of the way from the ventricle. In contrast, in *M. glutinosa*, the ventral aorta is unbranched and “I-shaped,” as in extant chondrichthyans (Marinelli & Strenger, 1956). This bifurcating structure in *E. burgeri* is identical to that of *E. stoutii* (Müller, 1834), but different from *E. cirrhatus* in which the ventral aorta bifurcates near the ventricle (Icardo et al., 2016a; see also Dean, 1904).

In contrast to the diversity in the branchial apparatus among the extant hagfish, all extant lampreys have seven bulsa branchialis and seven corresponding branchial pores, with less morphological diversity within the extant species than the hagfish. Although the morphology is uncertain in some extinct species, such as *†Hardistiella montanensis*, the Carboniferous *†Mayomyzon pieckoensis* has seven pairs of gills (Bardack, 1991; Miyashita et al., 2021 and references therein), indicating that the past diversity of lampreys might also have been much smaller than that of the extant hagfishes.

Similarly, whereas some hagfish species have a variable number of pharyngeal arches, all extant gnathostomes have a fixed number of pharyngeal arches. What allowed the deviation in the hagfish lineage from the developmental constraint that fixes the number of pharyngeal arches in other vertebrate taxa is unknown. It might have something to do with the development of the massive lingual apparatus, accompanied by a caudal shift of the branchial apparatus throughout development. Perhaps, the developmental process and time may have a redundancy in response to the extreme environment of the deep sea, and the number of gills might be affected by developmental adaptations to different marine environments.

In addition to their true heart (sometimes called the branchial heart), hagfish have multiple “accessory hearts” (Johansen, 1963; Nishiguchi et al., 2016). The significance and function of these accessory hearts are unknown, although some authors have suggested that they enable the storage of more than 30% of their blood in the sinus system to keep blood pressure low and ensure gill function (Forster, 1997). The number and locations of these accessory hearts are also known vary among species. For example, *M. glutinosa* has two cardinal hearts, one branchial heart, one portal heart, and one pair of caudal hearts, but *E. okinoseanus* does not have caudal hearts (Nishiguchi et al., 2016). In addition, hagfish maintain a lower blood pressure than other vertebrates, especially among the *Myxine* (Farrell, 2007). In hagfish living in deep-sea environments, the gill and cardiovascular systems appear to have been optimized for circulation at low blood pressure to maximize efficiency.

### Thyroid gland

The thyroid gland of the cyclostomes is not a glandular organ with clusters of follicles, but rather, the thyroid follicles are scattered throughout the pharynx (Kobayashi, 1987). Hagfish thyroid follicles are similar in early development and morphology to those of gnathostomes and represent the ancestral condition among extant vertebrates (Takagi et al., 2022). Electron microscopic analyses have revealed the presence of dense granules in the thyroid follicular epithelial cells of hagfishes, which are thought to contain iodine (Henderson & Gorbman, 1971; Fujita, 1975; Fujita & Shinkawa, 1975; Suzuki, 1985). Also, numerous microvilli are present on the apical membrane on the medial side of the follicular structure of thyroid follicular epithelial cells [> Also, numerous microvilli are present on the apical (luminal) membrane of thyroid follicular epithelial cells] (Suzuki and Kawabata, 1988). These observations indicate that the thyroid follicles of hagfish do not differ significantly from those of other vertebrates in terms of histology.

In lampreys, the thyroid gland takes on the function of an exocrine organ, known as an endostyle, during the larval period. This was long considered to be an ancestral condition, but recently it has been suggested to be a secondary condition (Takagi et al., 2022). Although there is some difference in the arrangement of blood vessels, both lamprey and hagfish have the scattered distribution of thyroid follicles along the bifurcated ventral aorta (for lamprey, see Takagi et al., 2022). This suggests that the ancestral thyroid gland was composed of scattered follicles, at least in the common ancestor of the cyclostomes, and that the thyroid as a glandular organ arose for the first time in the gnathostome lineage. A similar trend is seen in the pancreas, as discussed below.

### Hagfishes possess an ancestral-type hepatobiliary system

Morphological evolutionary studies of the hepatobiliary organs are less well-established than those of other body systems, such as the musculoskeletal system. Because of this, evolutionary studies of the hepatobiliary system are poorly developed, even though it is a characteristic structure of vertebrates.

The hepatobiliary system of the lamprey is distinct from that of jawed vertebrates, for example in having a single liver lobe (Marinelli & Strenger, 1954). Some older studies have claimed that the gallbladder is completely absent in lampreys, but this is incorrect; the gallbladder is present during the ammocoetes larval period and is lost in the adult stage (Scammon, 1916; Youson & Sidon, 1978; Youson, 1993; Morii et al., 2010). This degeneration of the gallbladder during metamorphosis is unique to lampreys. In this respect, adult lampreys are more specialized than their ammocoetes larvae.

In contrast to lampreys, the hepatobiliary organs of hagfish resemble those of jawed vertebrates at the gross anatomical level. For example, hagfish maintain two liver lobes and a large gallbladder throughout their lives (Fig. 5). Because of their elongated shape, the liver lobes are generally called anterior and posterior lobes, which correspond to the right and left lobes, respectively, of other vertebrates. The gallbladder is supplied by vessels branching from the artery of the body cavity (a. hepatica). The major veins of the liver gather in the vena portaea and enter the heart through a short route through the sinus venosus. The topographical relationships among the major veins are identical to those of mammals (Higashiyama et al., 2016) and birds (Higashiyama & Kanai, 2021), but distinct from those of lampreys. Thus, the hepatobiliary morphology is highly conserved among the vertebrates, and the lamprey case is likely to be a derived form.

Histologically, the hepatobiliary system of hagfish resembles that of jawed vertebrates, although with some important differences. Namely, in both hagfish and jawed vertebrates, the intrahepatic bile ducts and hepatic artery run parallel to the portal vein to form portal triads, and smooth muscle is distributed around the hepatic artery and gallbladder (Umezu et al., 2012; Shiojiri et al., 2019; Ota et al., 2021). However, it has recently been reported that in hagfish, both the intrahepatic and extrahepatic bile ducts and the portal vein lack smooth muscle layers. These characteristics have not been found in jawed vertebrates as far as is known (Ota et al., 2021).

Whereas lamprey and hagfish differ in the structure of their liver and bile ducts, both taxa are characterized by their lack of a compact pancreas (pancreatic argan). In extant mammals, the pancreas develops from the subdivision of a single embryonic primordium (the hepatic diverticulum) (Higashiyama et al., 2018), but this primordium is probably not present in cyclostomes. The exo- and endocrine cells exist sparsely in the intestinal wall (Yui et al., 1988; Youson & Al-Mahrouki, 1999), and could not be identified in our examination of gross anatomy. Thus, as with the thyroid, the pancreas as an organ as well as the pancreatic duct likely arose in the common ancestor of the jawed vertebrates, although we cannot exclude the possibility that the compact pancreas was lost secondarily in the common ancestor of the cyclostomes. Detailed embryological data is needed to provide further insight into this question.

## Conclusion

We dissected specimens of *E. burgeri* and compared its gross morphology, mainly around the pharynx, with that of other species. We identified several morphological differences between *E. burgeri* and other species in the pharyngeal region and the cardiovascular system but were not able to do so for other organs.

We did not conduct a detailed dissection of the musculoskeletal or nervous systems. Some differences in these systems could be tied to differences in ecology among hagfish species. For example, the eyes of *Myxine* are almost completely buried beneath the cutaneous layer (Marinelli & Strenger, 1956), whereas the position of the eyes can be identified externally in *Eptatretus*. It is plausible that there are differences in the shape of the brain and peripheral nerves associated with these sensory-organ differences. Comparison of the two sub-lineages will allow for a better understanding of hagfish evolution and development.

In the present study, we examined the anatomy of unfixed and fixed hagfish specimens. Although fixed specimens allow detailed observation of various body structures, the texture and color of each tissue are lost. In addition, sampling of the soft tissues, such as the thyroid gland and brain, is often performed under unfixed conditions. Therefore, the comparative dissection of unfixed and fixed specimens, herein described with photographs, should serve as a guide for future studies.

## Acknowledgments

We appreciate Hisashi Hasegawa and Kazutaka Hasegawa (Choukane Maru; Yaizu City, Shizuoka, Japan) providing the *E. atami* specimens and Hisanori Kohtsuka (Misaki Marine Station, the University of Tokyo) and Susumu Hatanaka (Shinsho Maru; Fujisawa, Kanagawa, Japan) providing the *E. burgeri* specimens. We are grateful to Kohei Kotake (Graduate School of Science, Shizuoka University) and Kazuma Nishio (Graduate School of Science, Shizuoka University) for their assistance in the dissection, and Kanako Sugawara (Graduate School of Science, Kyoto University) for providing the photograph of *E. burgeri* with exposed teeth in Figure 2. We also thank Dr. Hiroki Gotoh (Faculty of Science, Shizuoka University) for bringing us together and giving us the opportunity to write this research paper.

## Author contributions

B.M. designed research; B.M., D.G.S., and H.H. performed research; B.M., D.G.S., M.S., and H.H. analyzed data; B.M. and H.H. write the draft manuscript; and B.M., D.G.S., M.S., and H.H. reviewed and edited the manuscript.

## References

Bardack, D. (1991). First fossil hagfish (Myxinoidea): a record from the Pennsylvanian of Illinois. Science 254, 701–703.

Clark, A.J., & Summers, A.P. (2007) Morphology and kinematics of feeding in hagfish: possible functional advantages of jaws. J Exp Biol 210, 2897–3909.

Cole, F.J. (1907). XXVI.—A monograph on the general morphology of the myxinoid fishes, based on a study of *Myxine*. part II. The anatomy of the muscles. Earth and Environmental Science Transactions of The Royal Society of Edinburgh 45, 683–757.

Dean, B. (1904). Notes on Japanese Myxinoids: a new genus *Paramyxine*, and a new species *Homea okinoseana*, reference also to their eggs. The journal of the College of Science, Imperial University of Tokyo, Japan 19, 1–23.

Denisenko, E., Guo, B.B., Jones, M., Hou, R., De Kock, L., Lassmann, T., Poppe, D., Clément, O., Simmons, R.K., Lister, R., & Forrest, A.R. (2020). Systematic assessment of tissue dissociation and storage biases in single-cell and single-nucleus RNA-seq workflows. Genome Biology 21, 1–25.

Dong, E.M., & Allison, W.T. (2021). Vertebrate features revealed in the rudimentary eye of the Pacific hagfish (Eptatretus stoutii). Proc Biol Sci 288, 20202187.

Dupret, Y., Sanchez, S., Goujet, D., Tafforeau, P., & Ahlberg, P.E. (2014). A primitive placoderm sheds light on the origin of the jawed vertebrate face. Nature 507, 500–503.

Farrell, A.P. (2007). Cardiovascular systems in primitive fishes. Fish Physiology 26, 53–120.

Forster, M.E. (1997). The blood sinus system of hagfish: its significance in a low-pressure circulation. Comparative Biochemistry and Physiology Part A: Physiology 116, 239–244.

Fricke, R., Eschmeyer, W.N., & Van der Laan, R. (eds) (2022). Eschmeyer’s catalog of fishes: genera, species, references. (http://researcharchive.calacademy.org/research/ichthyology/catalog/fishcatmain.asp). Electronic version accessed 25 October 2022.

Fujita, H. (1975). X-ray microanalysis on the thyroid follicle of the hagfish, *Eptatretus burgeri* and lamprey, Lampetra japonica. Histochemistry 43, 283–290.

Fujita, H., Shinkawa, Y. (1975). Electron microscopic studies on the thyroid gland of the hagfish, *Eptatretus burgeri*. (a part of phylogenetic studies on the thyroid gland). Arch Histol Jap 37, 277–289.

Gabbott, S.E., Donoghue, P.C.J., Sansom, R.S., Vinther, J., Dolocan, A., & Purnell, M.A. (2016). Pigmented anatomy in Carboniferous cyclostomes and the evolution of the vertebrate eye. Proc R Soc B. 283, 20161151.

Gorbman, A., Kobayashi, H., Honma, Y., & Matsuyama, M. (1990). The hagfishery of Japan. Fisheries 15, 12–18.

Heimberg, A. M., Cowper-Sal· lari, R., Sémon, M., Donoghue, P. C., & Peterson, K. J. (2010). microRNAs reveal the interrelationships of hagfish, lampreys, and gnathostomes and the nature of the ancestral vertebrate. Proceedings of the National Academy of Sciences 107(45), 19379–19383.

Henderson, N.E., Gorbman, A. (1971). Fine structure of the thyroid follicle of the Pacific hagfish, Eptatretus stouti. Gen Comp Endocrinol 16, 409–429.

Higashiyama, H., Sumitomo, H., Ozawa, A., Igarashi, H., Tsunekawa, N., Kurohmaru, M., & Kanai, Y. (2016). Anatomy of the murine hepatobiliary system: A whole-organ-level analysis using a transparency method. Anat Rec (Hoboken) 299, 161–172.

Higashiyama, H., Uemura, M., Igarashi, H., Kurohmaru, M., Kanai-Azuma, M., & Kanai, Y. (2018). Anatomy and development of the extrahepatic biliary system in mouse and rat: a perspective on the evolutionary loss of the gallbladder. J Anat 232, 134–145.

Higashiyama, H., & Kanai, Y. (2021). Comparative anatomy of the hepatobiliary systems in quail and pigeon, with a perspective for the gallbladder-loss. J Vet Med Sci 83, 855–862.

Higuchi, S., Sugahara, F., Pascual-Anaya, J., Takagi, W., Oisi, Y., & Kuratani, S. (2019). Inner ear development in cyclostomes and evolution of the vertebrate semicircular canals. Nature 565, 347–350.

Hirasawa, T., Fujimoto, S., & Kuratani, S. (2016). Expansion of the neck reconstituted the shoulder–diaphragm in amniote evolution. Development, Growth & Differentiation 58, 143–153.

Honma, Y. (1998). Asian hagfishes and their fisheries biology. In The biology of hagfishes (pp. 45–56). Springer, Dordrecht.

Icardo, J.M., Colvee, E., Schorno, S., Lauriano, E.R., Fudge, D.S., Glover, C.N., & Zaccone, G. (2016a). Morphological analysis of the hagfish heart. I. The ventricle, the arterial connection and the ventral aorta. J Morphol 277, 326–340.

Icardo, J.M., Colvee, E., Schorno, S., Lauriano, E.R., Fudge, D.S., Glover, C.N., & Zaccone, G. (2016b). Morphological analysis of the hagfish heart. II. The venous pole and the pericardium. J Morphol 277, 853–865.

Janvier P. (1996). Early Vertebrates. Oxford: Oxford University Press pp. 393

Johansen, K. (1963). The cardiovascular system of *Myxine glutinosa*. In: Brodal A, Fänge R, editors. The biology of Myxine, Oslo: Universitetsforlaget. p 289–316.

Jørgensen, J.M., Lomholt, J.P., Weber, R.E., & Malte, H. (1998). The Biology of Hagfishes, Springer Dordrecht, pp 34–35.

Kase, M., Shimizu, T., Kamino, K., Umetsu, K., Sugiyama, H., & Kitano, T. (2017). Brown hagfish from the northwest and east coasts of Honshu, Japan are genetically different. Genes Genet Syst 92, 197–203.

Kuraku, S., Hoshiyama, D., Katoh, K., Suga, H., & Miyata, T. (1999). Monophyly of lampreys and hagfishes supported by nuclear DNA-coded genes. Journal of Molecular Evolution 49, 729–735.

Kuraku, S., & Kuratani, S. (2006). Time scale for cyclostome evolution inferred with a phylogenetic diagnosis of hagfish and lamprey cDNA sequences. Zoological Science 23, 1053–1064.

Kuraku, S. (2008). Insights into cyclostome phylogenomics: pre-2R or post-2R. Zoological Science, 25, 960–968.

Kuratani, S., Horigome, N., & Hirano, S. (1999). Developmental morphology of the head mesoderm and reevaluation of segmental theories of the vertebrate head: evidence from embryos of an agnathan vertebrate, Lampetra japonica. Dev Biol 210, 381–400.

Li, P., Conley, A., Zhang, H., & Kim, H.L. (2014). Whole-transcriptome profiling of formalin-fixed, paraffin-embedded renal cell carcinoma by RNA-seq. BMC genomics 15, 1–9.

Mallatt, J., & Sullivan, J. (1998). 28S and 18S rDNA sequences support the monophyly of lampreys and hagfishes. Molecular Biology and Evolution 15, 1706–1718.

Marinelli, W., & Strenger, A. (1954). Vergleichende Anatomie und Morphologie der Wirbeltiere, Band 1. Lampetra fluviatilis. Franz Deuticke, Wien.

Marinelli, W., & Strenger, A. (1956). Vergleichende Anatomie und Morphologie der Wirbeltiere Band 2. Myxine glutinosa. Franz Deuticke, Wien.

Meilhac, S.M., & Buckingham, M.E. (2018). The deployment of cell lineages that form the mammalian heart. Nat Rev Cardiol 15, 705–724.

Mincarone, M.M., & McCosker, J.E. (2004). Eptatretus lakeside sp. nov., a new species of five-gilled hagfish (myxinidae) from the Galápagos islands. *Reprinted from PCAS* 55, 162–168.

Miyashita, T., Coates, M.I., Farrar, R., Larson, P., Manning, P.L., Wogelius R.A., Edwards, N.P., Anné, J., Bergmann, U., Palmer, A.R., & Currie, P.J. (2019). Hagfish from the Cretaceous Tethys Sea and a reconciliation of the morphological–molecular conflict in early vertebrate phylogeny. PNAS 116, 2146–2151.

Miyashita, T., Gess, R.W., Tietjen, K., & Coates, M.I. (2021). Non-ammocoete larvae of Palaeozoic stem lampreys. Nature 591, 408–412.

Mizukami, K., Higashiyama, H., Arima, Y., Ando, K., Fukuhara, S., Miyagawa-Tomita, S., & Kurihara, H. (2022). Coronary artery established through amniote evolution. bioRxiv. doi: https://doi.org/10.1101/2022.09.06.506796

Morii, M., Mezaki, Y., Yamaguchi, N., Yoshikawa, K., Miura, M., Imai, K., Yoshino, H., Hebiguchi, T., & Senoo, H. (2010). Onset of apoptosis in the cystic duct during metamorphosis of a Japanese lamprey, Lethenteron reissneri. Anat Rec 293, 1155–1166.

Müller, J. (1834). Vergleichende Anatomie der Myxinoiden, der Cyclostomen mit durchbortem gaumen. I. Osteologie und myologie. Abhandlungen der königlichen Akademie der Wissenschaften zu Berlin. Wiss Berlin. pp. 65–340.

Nakabō, T. (Ed.). (2013). Fishes of Japan: with pictorial keys to the species, third edition. Tokyo: Tokai University Press.

Nelson, J.S., Grande, T.C., & Wilson, M.V. (2016). Fishes of the World. John Wiley & Sons.

Nishiguchi, Y., Tomita, T., Sato, K., Yanagisawa, M., Murakumo, K., Kamisako, H., Kaneko, A., Hiruta, N., Terai, K., Takahara, A., & Okada, M. (2016). Examination of the hearts and blood vascular system of *Eptatretus okinoseanus* using computed tomography images, diagnostic sonography, and histology. Int J Anal Bio-Sci 4, 46–54.

Nozaki, M., Ichikawa, T., Tsuneki K., & Kobayashi, H. (2000). Seasonal development of gonads of the hagfish, *Eptatretus burgeri*, correlated with their seasonal migration. Zoolog Sci 17, 225–232.

Oisi, Y., Ota, K.G., Kuraku, S., Fujimoto, S., & Kuratani, S. (2013). Craniofacial development of hagfishes and the evolution of vertebrates. Nature 493, 175–180.

Oisi, Y., Fujimoto, S., Ota, K.G., & Kuratani, S. (2015). On the peculiar morphology and development of the hypoglossal, glossopharyngeal and vagus nerves and hypobranchial muscles in the hagfish. Zoological Letters 1, 1–15.

Ota, K.G., Kuraku, S., & Kuratani, S. (2007). Hagfish embryology with reference to the evolution of the neural crest. Nature 446, 672–675.

Ota, K.G., Fujimoto, S., Oisi, Y., & Kuratani, S. (2011). Identification of vertebra-like elements and their possible differentiation from sclerotomes in the hagfish. Nat Commun 2, 373.

Ota, N., Hirose, H., Kato, H., Maeda, H., & Shiojiri, N. (2021). Immunohistological analysis on distribution of smooth muscle tissues in livers of various vertebrates with attention to different liver architectures. Annals of Anatomy-Anatomischer Anzeiger 233, 151594.

Patzner, R.A. (1978a). Cyclical changes in the testis of the hagfish *Eptatretus burgeri* (Cyclostomata). Acta Zool. 58, 223–226.

Patzner, R.A. (1978b). Cyclical changes in the ovary of the hagfish *Eptatretus burgeri* (Cyclostomata). Acta Zool. 59, 57–61.

Romagnani, P., Lasagni, L., & Remuzzi, G. (2013). Renal progenitors: an evolutionary conserved strategy for kidney regeneration. Nat Rev Nephrol 9, 137–146.

Scammon R.E. (1916). On the development of the biliary system in animals lacking a gallbladder in postnatal life. The Anatomical Record 10, 543–558.

Shiojiri, N., Tanaka, S., & Kawakami, H. (2019). The hepatic architecture of the coelacanth differs from that of the lungfish in portal triad formation. Okajimas Folia Anatomica Japonica 96, 1–11.

Sugahara, F., Murakami, Y., Pascual-Anaya, J., & Kuratani, S. (2017). Reconstructing the ancestral vertebrate brain. Dev Growth Differ 59, 163–174.

Sugahara, F. (2021). Cyclostomes (Lamprey and Hagfish). In: Schierwater, B. & Boutet, A., eds. Handbook of Marine Model Organisms in Experimental Biology. CRC Press: Boca Raton. pp. 403–417.

Suzuki, D.G. (2021). Consciousness in jawless fishes. Frontiers in Systems Neuroscience 15, 751876.

Suzuki, S. (1985). Iodine distribution in the thyroid follicles of the hagfish, *Eptatretus burgeri* and lamprey, *Lampetra japonica:* Electron-probe X-ray microanalysis. Cell Tissue Res 241, 539–543.

Suzuki, S. (1992). 4 Thyroid gland. In: Matsumoto, A., Ishii, S., eds. Atlas of Endocrine Organs: Vertebrates and Invertebrates, Springer: Berlin. pp. 63–71.

Suzuki, S., & Kawabata, I. (1988). A scanning electron microscopic study on the thyroid follicle of the hagfish Eptatretus burgeri. Acta Zool 69, 253–258.

Takagi, W., Sugahara, F., Higuchi, S., Kusakabe, R., Pascual-Anaya, J., Sato, I., Oisi, Y., Ogawa, N., Miyanishi, H., Adachi, N., Hyodo, S., & Kuratani, S. (2022). Thyroid and endostyle development in cyclostomes provides new insights into the evolutionary history of vertebrates. BMC biology 20, 1–10.

Takezaki, N., Figueroa, F., Zaleska-Rutczynska, Z., & Klein, J. (2003). Molecular phylogeny of early vertebrates: monophyly of the agnathans as revealed by sequences of 35 genes. Molecular Biology and Evolution 20, 287–292.

Tzahor, E., & Evans, S.M. (2011). Pharyngeal mesoderm development during embryogenesis: implications for both heart and head myogenesis. Cardiovasc Res 91, 196–202.

Umezu, A., Kametani, H., Akai, Y., Koike, T., & Shiojiri, N. (2012). Histochemical analyses of hepatic architecture of the hagfish with special attention to periportal biliary structures. Zoolog Sci 29, 450–457.

Yalden, D.W. (1985). Feeding mechanisms as evidence for cyclostome monophyly. Zoological Journal of the Linnean Society 84, 291–300.

Yamatsuta, K. (1903). The anatomy of *Bdellostoma cirrhatum*, Gthr. Bulletin of Tokyo Kōtō Shihan Gakkō Hakubutsu Gakkai 1, 1–3. (in Japanese)

Yokoyama, H., Yoshimura, M., Suzuki, D.G., Higashiyama, H. & Wada, H. (2021). Development of the lamprey velum and implications for the evolution of the vertebrate jaw. Dev Dyn 250, 88–98.

Youson, J., & Sidon, E. (1978). Lamprey biliary atresia: First model system for the human condition? Cellular and Molecular Life Sciences 34, 1084–1086.

Youson, J.H. (1993). Biliary atresia in lampreys. Adv Vet Sci Comp Med 37, 197–255.

Youson, J.H., & Al-Mahrouki, A.A. (1999). Ontogenetic and phylogenetic development of the endocrine pancreas (islet organ) in fish. Gen Comp Endocrinol 116, 303–335.

Yui, R., Nagata, Y., & Fujita, T. (1988). Immunocytochemical studies on the islet and the gut of the arctic lamprey, Lampetra japonica. Arch Histol Cytol 51, 109–119.

Ziermann, J.M., Miyashita, T., & Diogo, R. (2014). Cephalic muscles of Cyclostomes (hagfishes and lampreys) and Chondrichthyes (sharks, rays and holocephalans): comparative anatomy and early evolution of the vertebrate head muscles. Zool J Linn Soc-Lond 172, 771–802.

